# Copepod functional traits and groups show divergent biogeographies in the global ocean

**DOI:** 10.1101/2022.02.24.481747

**Authors:** Fabio Benedetti, Jonas Wydler, Meike Vogt

## Abstract

**Aim:** The distribution of zooplankton functional traits is a key factor for regulating food web dynamics and carbon cycling in the oceans. Yet, we lack a clear understanding of how many functional groups (FGs) exist in the zooplankton and how their traits are distributed on a global scale. Here, we model and map the environmental habitats of copepod (i.e. the main component of marine zooplankton) FGs to identify regions sharing similar functional trait expression, at the community level.

**Taxon:** Marine planktonic Neocopepoda.

**Location:** Global ocean.

**Methods:** Factor analysis on mixed data and hierarchical clustering were used to identify copepod FGs based on five species-level functional traits. An ensemble of species distribution models was used to estimate the environmental niches of the species modelled and the community weighted mean values of the traits studied. Ocean regions were defined based on their community-level mean trait expression using a principal component analysis and hierarchical clustering.

**Results:** Eleven global copepod FGs were identified. They displayed contrasting latitudinal patterns in mean annual habitat suitability that could be explained by differences in environmental niche preferences: two FGs were associated with polar conditions, one followed the global temperature gradient, five were associated with tropical oligotrophic gyres, and the remaining three with boundary currents and counter currents. Four main regions of varying community weighted mean trait values emerged: the Southern Ocean, the northern and southern high latitudes, the tropical gyres, and the boundary currents and upwelling systems.

**Conclusions:** The present FGs will improve the representation of copepods in global marine ecosystem models. This study improves the understanding of the patterns and drivers of copepods trait biogeography and will serve as a basis for studying links between zooplankton biodiversity and ecosystem functioning in a context of climate change.

## 1. Introduction

Copepods are crustaceans that dominate the biomass of the mesozooplankton (0.2-2.0 mm) and rank amongst the most abundant animals in the oceans (Huys & Boxshall, 1991). They are morphologically and functionally diverse and adapted to all aquatic ecosystems, with particular success in marine systems (Huys & Boxshall, 1991; Kiørboe, 2011a). Copepods play a pivotal role in food webs, both as microplankton grazers and prey for higher trophic levels (Beaugrand, Edwards, & Legendre, 2010). Copepods also play a critical role in the marine biological carbon pump by performing extensive vertical migrations, grazing on phytoplankton in the sunlight layers and releasing relatively fast-sinking fecal pellets, exporting organic carbon with dead carcasses down to the deep layers (Turner, 2015; Steinberg & Landry, 2017).

The relative contribution of planktonic copepods to the above-mentioned processes is mediated by their diversity and trait expression (Barton et al., 2013). For instance, larger copepods perform stronger diel vertical migrations and produce larger fecal pellets that sink faster and thus increase the proportion of carbon exported to depth (Stamieszkin et al., 2015; Ohman & Romagnan, 2016). The dominance of certain copepod feeding modes impacts food-web dynamics as feeding modes affect grazing and mortality rates (Kenitz, Visser, Mariani, & Andersen, 2017; van Someren Gréve, Almeda, & Kiørboe, 2017). Yet, mechanistic ecosystem models usually represent zooplankton only through a few size classes (Le Quéré et al., 2005), which oversimplifies the contribution of species and functional diversity to ecosystem functioning and biogeochemical cycles (Flynn et al., 2015). Multiple studies show that the number and grazing characteristics of zooplankton functional groups (FGs) control the biomass and diversity of other functional groups, with consequences for global biogeochemical cycles (Prowe et al., 2012; Sailley et al., 2013; Vallina et al., 2014; Le Quéré et al., 2016). Ecosystem models rely strongly on parameterisations of traits governing biological processes and food-web interactions based on scarce empirical evidence (Barton et al., 2013). Consequently, investigating the potential links between copepod trait distribution and ecosystem functioning contributes to improving these models (Stocker, 2014).

To better represent the role of zooplankton in marine systems, ecologists are increasingly adopting trait-based approaches (Litchman, Ohman, & Kiørboe, 2013; Hébert, Beisner, & Maranger, 2017). Functional traits are species- or organism-level characteristics that affect their fitness and are linked to survival, feeding, growth, and reproduction (Violle et al., 2007; Litchman et al. 2013). Due to trade-offs between them, organisms cannot maximize all functions at the same time. For example, copepods that rely on active ambush feeding (i.e., copepods that only move to capture prey) show 8.5 times lower mortality rates than copepods that rely on active feeding strategies (van Someren Gréve, Almeda, & Kiørboe, 2017). Yet, adult male copepods need to move actively to find a mate for reproduction, which undermines the benefits of their feeding mode (Kiørboe, 2011b; Litchman et al., 2013).

Copepod traits dynamics can be investigated by grouping species into FGs with similar trait combinations (Benedetti, Gasparini, & Ayata, 2016; Becker et al., 2021). Categorizing species according to their similarity in functional traits rather than their taxonomic classification enables to summarize their diversity into distinct, parsimonious groups. At the regional scale, Pomerleau et al. (2015) showed that 42 zooplankton species in the North East Pacific Ocean could be clustered into five groups based on their body length, feeding and reproduction mode, and trophic group. Similarly, Benedetti, Vogt et al. (2018) grouped 106 dominant Mediterranean copepod species into seven FGs with distinct environmental niche characteristics. Such groupings can improve the representation of zooplankton in ecosystem models since FGs increase the representation of ecological function without adding taxonomic diversity and complexity.

The functional composition of zooplankton can be examined using functional diversity indices which indicate how changes in species richness and composition affect the emergent expression of traits (Villéger, Mason & Mouillot, 2008). These indices allow to investigate biodiversity and ecosystem functioning relationships as they indicate changes in ecological processes that drive community assembly. For instance, Becker et al. (2021) documented a poleward decrease in copepod functional diversity in the South Atlantic, which was driven by the decreasing contribution of cruising carnivores and small and medium-sized broadcasting copepods. The authors attributed this poleward decrease in functional diversity to niche partitioning as only a subset of functionally very dissimilar groups of copepods thrived in the cold and productive waters of the Southern Ocean.

The functional composition of zooplankton can also be investigated through community weighted mean (CWM) values of traits (Ricotta, 2005; Brun et al., 2016), i.e. the proportion of species in a community that exhibits a particular trait value weighed by their relative contribution to community composition. Thereby, CWM values of functional traits help to identify the relationships between the emerging expression of functional traits on a community-level, the environmental drivers of their biogeographical patterns of trait expression, and the potential trade-offs underlying trait expression. The first truly global study by Brun et al. (2016) found larger copepod body sizes, higher proportions of myelinated species, and relatively smaller clutch size at latitudes >50°, associated with gradients in temperature and phytoplankton seasonality. To our knowledge, other qualitative traits such as trophic group or spawning mode have been investigated but not using a global CWM trait approach (McGinty et al., 2018; Benedetti, Vogt et al., 2018). If patterns of CWM values are clustered in space and time, they can be used to define ocean regions sharing common ecological characteristics (Longhurst, 2010; Reygondeau et al., 2017; Hofmann Elizondo et al., 2021). Although new data compilations make it possible, neither copepod FGs nor CMW-based regionalization schemes have been investigated on a global scale, yet this could be highly informative of large-scale marine food web and ecosystem dynamics.

Here, we use hierarchical clustering and a state-of-the-art species distribution models (SDMs) framework to address the following questions: (i) What are the global copepod FGs emerging from the clustering of the functional traits of commonly sampled copepod species? (ii) Do copepod FGs show different environmental niche preferences (i.e. niche centers and widths)? (iii) What are the habitat distribution patterns of these FGs and their community-level trait expression, and which regions of the global ocean share similar trait expression?

## 2. Materials and Methods

Figure 1 summarizes the analytical framework we developed to identify emergent copepod FGs and their habitats in the global ocean using an ensemble species distribution model (SDMs) approach.

**Figure 1:**
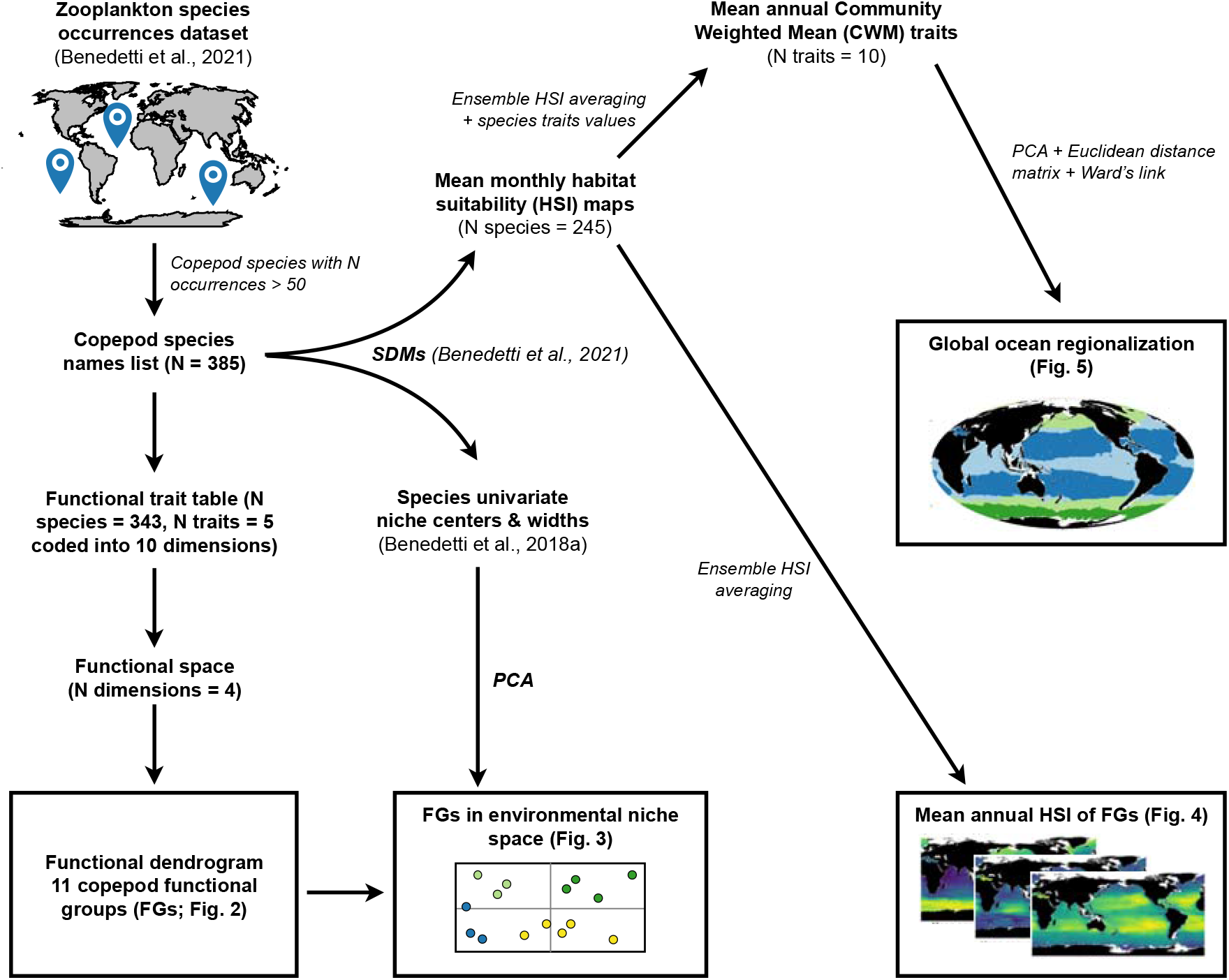
Flowchart summarizing the numerical analyses implemented in our study. SDMs = Species distribution models, PCA = Principal component analysis.

### 2.1. Species occurrence data

We used the zooplankton occurrences dataset (geolocated and dated presences) compiled by Benedetti et al. (2021). Occurrences corresponding to benthic and parasitic copepods, occurrences with missing spatial coordinates, sampling dates, sampling depth, or not identified at species-level were removed. Data from drilling holes, grid cells within 25 km of the nearest shoreline, or with a maximum sampling depth >500 m were also discarded (see Benedetti et al., 2021). To address sampling effort biases in the copepod occurrence data (Appendix S1) that could inflate models performance metrics or over-represent portions of the environmental space and thus hinder model predictability and interpretation (Veloz, 2009; Hijmans, 2012), we thinned occurrences using a randomization algorithm for each month and each species (30 randomizations per species dataset) so that monthly species occurrences were at least 500 km apart (Aiello-Lammens, Boria, Radosavljevic, Vilela, & Anderson, 2015). Only those 385 species displaying at least 50 occurrences were retained to constitute the list of species for which functional traits data were retrieved (Appendix S2).

### 2.2. Species functional trait data

Five functional traits were included based on data availability from the literature (Appendices S2 and S3):

Body size (quantitative continuous): mean maximum adult female body size (i.e. length of the cephalothorax) in mm. Body size is considered a master trait as it impacts all life functions, scales with most physiological rates and influences predator-prey interactions (Hansen, Bjornsen, & Hansen, 1994; Kiørboe & Hirst, 2014; Hébert, Beisner, & Maranger, 2017). The mean maximum body size values used here were derived from Razouls et al. (2015) and range from 0.46 to 11.00 mm.

Trophic group (categorical): most marine planktonic copepods are omnivorous but they can be grouped according to their primary food source to describe their role in food-web dynamics (Pomerleau et al., 2015; Benedetti et al., 2016). Here, the following five groups were defined: Omnivore-Herbivore, Omnivore-Carnivore, Omnivore-Detritivore, strict Carnivore, and Omnivore.

Feeding mode (categorical): copepods rely on various strategies to detect and capture their prey. We follow the definitions of Kiørboe (2011a): ambush-feeding, current-feeding, cruise-feeding, particle-feeding, current-cruise feeding, and current-ambush feeding (the last two refer to mixed-feeding species with both strategies). Ambush-feeding copepods lurk in the water column, detect the vibration generated by motile preys and capture them through quick jumps. Current-feeding species use a scanning current to detect and capture immobile preys like phytoplankton cells (Kiørboe, 2011a). Cruise-feeders are copepods that swim actively through the water column in search of their prey. Lastly, particle-feeders are species known to live on aggregates of sinking organic particles such as discarded appendicularians’ houses. Myelination (binary): myelinated copepod species have a lipid-rich myelin sheath around their nerves that enables faster attack or evasive reactions (Lenz, 2012). Myelin sheaths play a key role in modulating mortality and feeding rates and improve energy savings at low food conditions.

Spawning mode (binary): eggs are either released into the open water after fertilization (free-spawning) or are carried by females in egg sacs or in egg masses (sac-spawning). Sac spawning copepods display lower fecundity rates and longer hatching times (Kiørboe & Sabatini, 1994).

### 2.3. Definition of functional groups

Only those copepod species with a body size value and no missing data for three out of the five traits were retained (n = 343). The species’ trophic groups and feeding modes were recoded binarily to accurately represent species that belong to several feeding modes/ trophic groups (Appendix S2). Thus, the analyses described below were performed on 343 species based on ten functional trait dimensions.

A factor analysis on mixed data (FAMD; Pagès, 2004) that summarize the main modes of trait combinations was used to estimate the functional distances between species. The use of a FAMD is an improvement to the Multiple Correspondence Analysis used in Benedetti et al. (2016) as it allowed to retain body size as a quantitative continuous variable. Four FAMD components explaining 80.15% of the variance in species traits were retained (Audigier et al., 2016). A Euclidean distance matrix was computed from the species’ score along the four FAMD components, and Ward’s agglomeration link was used to generate a functional dendrogram. The number FGs was determined by the height at which the dendrogram is cut. We investigated the sensitivity of the resulting FGs to the main parameters of our clustering approach (Appendix S4) and inspected each dendrogram to ensure the ecological relevance of the final FGs (i.e., avoid few large groups that are functionally heterogenous or numerous small groups that are functionally redundant). Ultimately, eleven FGs were identified (Table S4.2).

### 2.4. Species distribution modelling

#### 2.4.1. Environmental predictor selection

We considered 14 commonly used environmental drivers of the spatial ranges of marine copepod species (see Appendix S5; Benedetti et al., 2021). The monthly climatologies of all predictor variables were projected onto the 1°x1° cell grid of the World Ocean Atlas (WOA; Boyer et al., 2013) and matched with the species occurrence data.

We used a two-stage procedure to select environmental predictors. First, we removed collinear predictors (mean Spearman rank correlation coefficient > |0.7|) to avoid increasing the uncertainty in regression model projections through coefficient inflation (Dormann et al., 2013). This first step retained the following ten predictors: sea surface temperature (SST, °C), photosynthetically active radiation (PAR, μmol m^-2^ s^-1^), logged nitrate concentration (logNO_3_, μM), mixed layer depth (MLD, m), logged chlorophyll a concentration (logChla, mg m^-3^), logged eddy kinetic energy (logEKE, m^2^ s^-2^), surface wind speed (Wind, m s^-1^), surface carbon dioxide partial pressure (pCO_2_, μatm), the excess of silicates to nitrates (Si^*^ = [SiO_2_] - [NO_3_^-^], μM) and the excess of nitrate over phosphate (N^*^ = [NO_3_] - 16[PO_4_^3-^], μM).

Second, the number of environmental predictors included in the SDMs was restricted to five to achieve a 10:1 ratio of occurrences to predictors (Guisan et al., 2017). For each species and SDM, one of the ten predictors was randomly reshuffled prior to training a model (n = 30 repetitions). The correlation coefficients between the predictions of the original dataset and those of the reshuffled datasets were used to estimate the relative ranking of each predictor (Appendix S6). We used the top five predictors for each FG and SDM separately to train the SDMs and project the species’ habitat suitability (Appendix S6).

#### 2.4.2. Background data, model configuration and habitat suitability projections

Pseudo-absence data were generated to use correlative SDMs following the target-group approach of Phillips et al. (2009) which is appropriate to model plankton groups with sparse data (Righetti et al., 2019; Benedetti et al., 2021). We chose to randomly draw pseudo-absences from the total pool of sites (i.e., monthly 1° x 1° grid cells) with at least ten occurrences across all species, except for FG2, FG4 and FG11 which showed sampling efforts that were distinct from the other groups (Appendices S1 and S4). For species of FG2, FG4 and FG11, pseudo-absences were randomly drawn only from those sites where at least 10 different species of their FG were found. Ten times more background data than presences were generated and presences were weighted ten times more than pseudo-absences (Barbet-Massin et al., 2012).

We developed an ensemble modelling approach that combined three SDMs types which were configured to avoid model overfitting (Merow et al., 2014; Thuiller et al., 2020): Generalized Additive Models (GAMs), Generalized Linear Models (GLMs), and Artificial Neural Networks (ANNs). The GLM were run using a quadratic formula and a logit link function. The GAMs were set up to use a logit link function and a maximum of five smoothing terms. ANNs were run using five cross validations with 200 iterations to optimize number of unities in the hidden layer and the parameter for weight decay. The SDMs were trained on 80% of the presence/pseudo-absence data chosen at random and tested on the remaining 20%. Ten random cross evaluations were carried out and the true skill statistic (TSS) was calculated to evaluate model performance. Only species displaying a mean TSS score >0.30 were used for the final species and functional trait projections. The SDMs were then projected onto the monthly environmental predictor conditions of the global ocean to obtain species-level maps of habitat suitability indices (HSI). For each FG, annual mean HSI values were obtained by averaging the monthly HSI maps across each species constituting the FG and SDMs.

### 2.5. Positioning FGs in environmental niche space

The mean univariate response curves emerging from GAMs were used to estimate the relative optimal conditions (i.e., niche center) and tolerance (i.e., niche width) for each species and predictor (Appendix S7; Benedetti, Vogt et al., 2018). The niche center was calculated as the HSI-weighted median and the niche width as the difference between the weighted 10^th^ and 90^th^ percentiles. For this analysis, a set of eight predictors common to all species was defined based on the overall ranking of predictors (i.e., unlike the top five FG-specific predictors used for global SDMs projections; Appendix S6): SST, logChl, logNO_3_, MLD, Si^*^, PAR, N^*^ and logEKE. We do this because a principal component analysis (PCA; Legendre & Legendre, 2012) was then performed on the niche centers and niche widths of all eight common predictors to ordinate the species in an environmental “niche space”. Non-parametric variance analyses (Kruskal-Wallis tests and post-hoc Dunn’s test) were carried out to test if FGs differ in their position in niche space.

### 2.6. Community weighted median (CWM) proportions of traits

To explore the biogeography of all copepod traits, the monthly CWM trait values were computed as the weighted sum of all HSI values belonging to species exhibiting that trait over the sum of HSI values across all species in that community (Cormen et al., 2009). Like for FGs, monthly CWM projections were performed for every SDM and mean annual CWM trait values were derived based on the ensemble of projections (Appendices S8 and S9).

To explore the regional patterns of copepod traits expression, clustering was used to divide the global ocean into regions with similar mean annual projections of: CWM body size, trophic groups, feeding modes, myelination and spawning mode (Appendix S10). The spatial patterns of the twelve CWM traits values were summarized through a PCA and the scores of each grid cell along the first four components (97.9% of total variance, Appendix S10) were used to calculate an Euclidean distance matrix. Various clustering approaches were explored based on this distance matrix (Appendix S11). Ultimately, we chose to present the CWM trait patterns obtained for k = 4 under Ward’s linkage based on the profiles of cluster stability metrics and because this method minimizes intra-cluster variance.

## 3. Results

### 3.1. Copepod FGs

The 343 copepod species retained were clustered based on their combinations of body size, feeding mode, trophic group, myelination, and spawning mode. Eleven copepod FGs could be derived from the functional dendrogram (Fig. 2; Appendix S4). The first dichotomy in the dendrogram occurred between non carnivorous cruise feeders (FG1, 2, 3) and all other FGs. FG1 was composed of 14 cruise-feeding herbivores of rather small sizes (median ± IQR = 1.10 ± 0.52 mm). Almost all species (13 out of 14) were sac-spawning, myelinated and belonged to the *Clausocalanus* genera. FG2 was defined by small (0.80 ± 0.30 mm) cruise-feeding detritivores. The 22 species of this group were all amyelinated, sac-spawning and belonged to the Oncaeidae family. FG3 was composed of 46 detritivores spanning a broader ranges of sizes (2.61 ± 1.70 mm) than the two previous groups. Most species in FG3 were myelinated (74%) and free-spawning (80%). Cruise- and current-feeding were the two predominant feeding modes, with *Spinocalanus, Metridia* and *<u>Scaphocalanus</u>* being the dominant genera in this FG.

**Figure 2:**
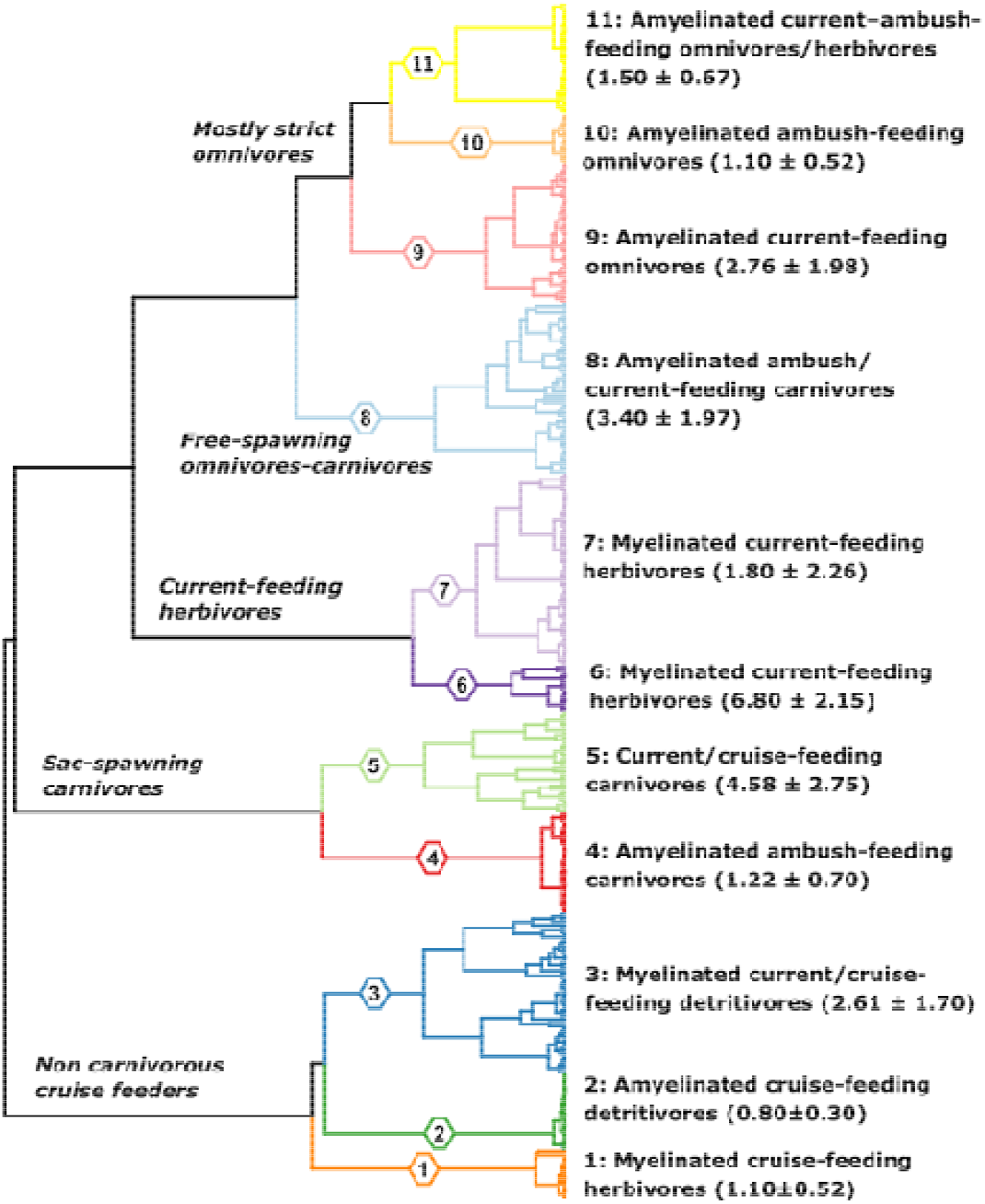
Functional dendrogram representing the inter-species traits dissimilarity for 343 copepod species. The hierarchical clustering was performed on a Euclidean distance matrix issued from the species coordinates ensuing from a Factor Analysis of Mixed Data (FAMD). Each leaf of the functional dendrogram represents a copepod species and the eleven functional groups (FGs) are numbered and highlighted in color. The range (median ± IQR) of the mean maximum body size of each FG is given in brackets.

The second dichotomy in the dendrogram separated the small and large sac-spawning carnivores (FG4 and 5) from the rest. FG4 consisted of 29 rather small (1.22 ± 0.70 mm) carnivorous, sac-spawning ayelinated ambush-feeders part of the Corycaeidae family and thus belonged to either the *Corycaeus, Farranula* or *Vettoria* genera. FG5 was made up of 29 of larger (4.58 ± 2.75 mm) species of sac-spawning current- and cruise-feeding carnivores from the Sapphirinidae and Euchaetidae families.

The third dichotomy of the dendrogram separated the large and medium-sized current feeders (FG6 and 7) from the rest. FG6 was a group of 13 rather large (6.80 ± 2.15 mm) myelinated, free-spawning species that were either current-feeding herbivores (85%) or fully omnivorous (15%). The two most represented genera were *Calanus* and *Eucalanus*. FG7 was the largest FG with 55 and current-feeding herbivores spanning a rather wide size range (1.80 ± 2.26 mm). Most were myelinated (95%) and free-spawning (89%). The main genera contributing to the composition of FG7 were the smaller *Calanus* (e.g., *C. helgolandicus, C. finmarchicus* or *C. pacificus*), and *Calocalanus* and *Paracalanus*.

The fourth dichotomy occurred between the small and medium-sized omnivores and the medium free spawning carnivores of FG8. FG8 consisted of 49 predominantly amyelinated (86%) and free-spawning (80%) medium-sized omnivorous-carnivorous species of various size (3.40 ± 1.7 mm). This diverse group mixed current- (60%) and ambush-feeders (40%) and gathered 12 genera with *Candacia, Haloptilus* and *Heterorhabdus* being the most dominant. FG9 was composed of 40 free-spawning and mostly amyelinated (72%). current-feeding omnivorous species showing a body sizes range of 2.76 ± 1.98 mm. *Pleuromamma, Gaetanus*, and *Labidocera* were the main genera.

Finally, the fifth and last main dichotomy separated the mixed-feeding omnivores (FG11) from the smaller ambush feeders (FG10). FG10 was rather homogeneous and contained 14 small (1.10 ± 0.52 mm) ambush-feeding, amyelinated and sac-spawning omnivores. All species belonged to either the *Oithona* or the *Dioithona* genera from the Oithonidae family. FG11 was a group of 32 amyelinated, free-spawning, current-ambush feeding species predominantly belonging to the *Acartia* (omnivorous-herbivorous) and *Centropages* (omnivorous) genera. All species in FG11 were rather small (1.50 ± 0.67 mm).

The functional space defined by the PCs of the FAMD describes the reduced functional space (Appendix S12). The first four PC of the FAMD explained 79.21% of the total variance in functional traits. The largest functional distance along the first component was found between FG4 and FG6.

### 3.2. FGs in environmental niche space

The univariate niche centers and widths of the copepod species modelled were used to position FGs in niche space (Fig. 3; Appendix S13). The main niche characteristics that contributed the most positively to PC1 (relative contribution to PC1 given in brackets) were: logChl center (12.16%), logChl width (12.51%), EKE width (11.78%), MLD center (9.96%), and MLD width (10.78%). The most important niche characteristics scoring PC2 were: Si^*^ center (30.88%), Si^*^ width (27.90%) and SST center (11.17%). Species with positive PC1 scores were those affiliated with wider niches and conditions of higher concentrations of nutrients and chlorophyll-a, stronger seasonal variations, and overall higher water column turbulence and mixing.

**Figure 3:**
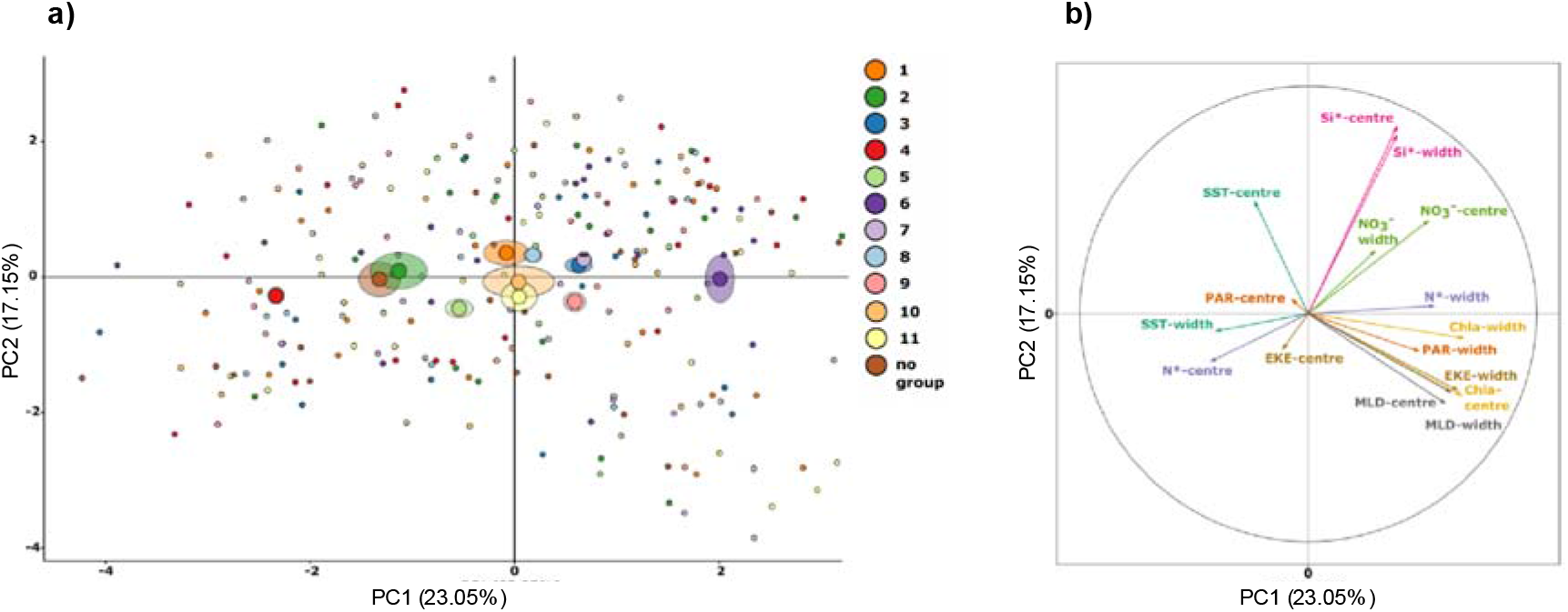
Position of a) functional groups (FGs, larger circles) and species (smaller circles) in environmental niche space according to the first two principal components (PCs) of a principal component analysis (PCA) performed on b) the species-level niche characteristics derived from GAM-based univariate response curves. The semi-transparent ellipses indicate two times the value of the standard errors associated with the mean PC scores of the FGs.

Post-hoc variance analyses showed significant (Dunn’s tests; p < 0.05) inter-FGs variations in niche characteristics and PC scores (Appendix S14). The largest distance in niche space (PC1) was found between FG4 and FG6 (Fig. 3a; p = 6.5e-10). Along PC1, FG4 also differed significantly from FG1 (p = 0.042), FG3 (p = 1.6e-06), FG7 (p = 1.3e-08), FG8 (p = 1.4e-4), FG9 (p = 2.3e-06), FG10 (p = 0.011) and FG11 (p = 0.005). Conversely, FG6 differed significantly from FG2 (p = 4.5e-4), FG5 (p = 0.001) and FG8 (p = 0.049) along PC1. FG6 also differed significantly from FG2 (p = 0.001) and FG5 (p = 0.002). None of the FGs showed significant variations along PC2. Only FG3 and FG7 showed significant variations along PC3 (p = 4.2e-3; Appendices S13 and S14). To summarize, when FGs 4 and 6 were not accounted for, none of the remaining nine FGs showed significant variations in PC1 and PC2 scores (all p > 0.05).

### 3.3. Mean annual habitat suitability indices (HSI) patterns

For all three SDMs and 11 FGs combined, we found SST to be the most important predictor, followed by PAR, logNO_3_, MLD, logChl, logEKE, Si^*^, N^*^, Wind, and pCO_2_ (Appendix S6). Thus, most FGs displayed mean annual HSI patterns that were driven by the latitudinal temperature gradient at the first-order (Fig. 4).

**Figure 4:**
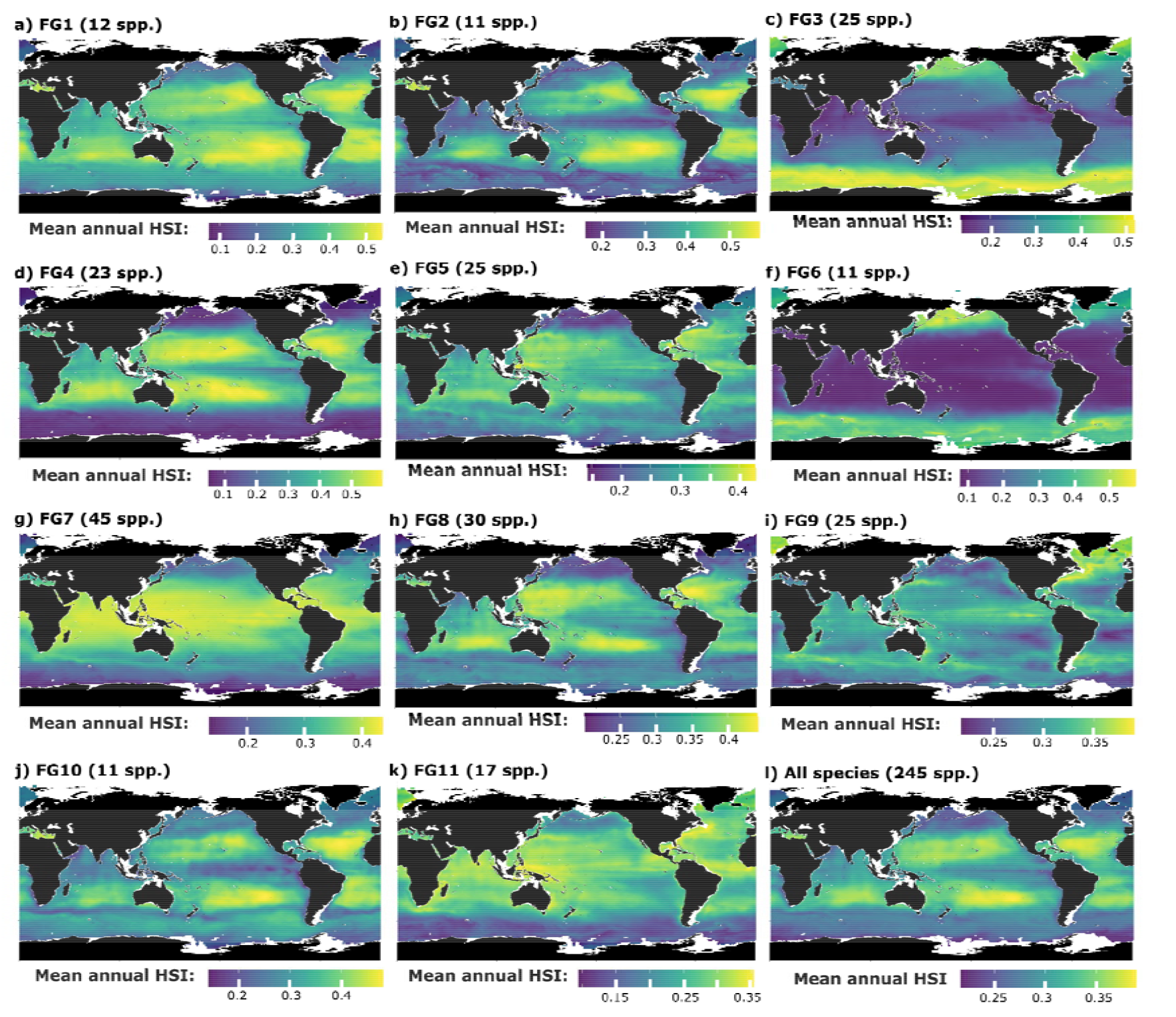
Mean annual habitat suitability index (HSI) of the eleven copepod functional groups (FG) for the global surface ocean (Mercator projection): a) FG1 (myelinated cruise-feeding omnivores-herbivores), b) FG2 (amyelinated cruise-feeding detritivores), c) FG3 (myelinated mixed-feeding or cuise-feeding detritivores), d) FG4 (amyelinated ambush-feeding carnivores), e) FG5 (current- or cruise-feeding carnivores), f) FG6 (myelinated current-feeding omnivores-herbivores), g) FG7 (myelinated current-feeding omnivores-herbivores), h) FG8 (amyelinated ambush- or current-feeding carnivores), i) FG9 (amyelinated current-feeding omnivores), j) FG10 (amyelinated ambush-feeding omnivores), k) FG11 (amyelinated mixed-feeding omnivores) and for l) all species together. Mean annual estimates were derived from the 12 monthly estimates of mean HSI obtained for three species distribution models (SDMs).

The mean annual HSI of FG3 was found to decrease progressively from the poles towards the equator. In contrast, the mean annual HSI of FG1, FG2, FG4, FG7, FG8, and FG10 were found to be highest in the tropics and decreased towards higher latitudes. For those five FGs, slighter decreases in HSI were modelled towards the tropical upwelling systems (e.g., Peru, Benguela), the northern part of the Indian Ocean and the Pacific Equatorial counter current. Therefore, these five FGs reached maximal HSI in the oligotrophic conditions of the tropical gyres. Meanwhile, FG5, FG9, and FG11 show less marked latitudinal gradients in mean annual HSI values, but they all displayed lower HSI in higher latitudes than in the tropics.

### 3.4. Functional trait biogeography from CMW trait values and regionalization

We identified four ocean regions that displayed significant contrasts in CWM trait proportions and that overlap with major oceanographic features (Fig. 5; Appendix S15).

**Figure 5:**
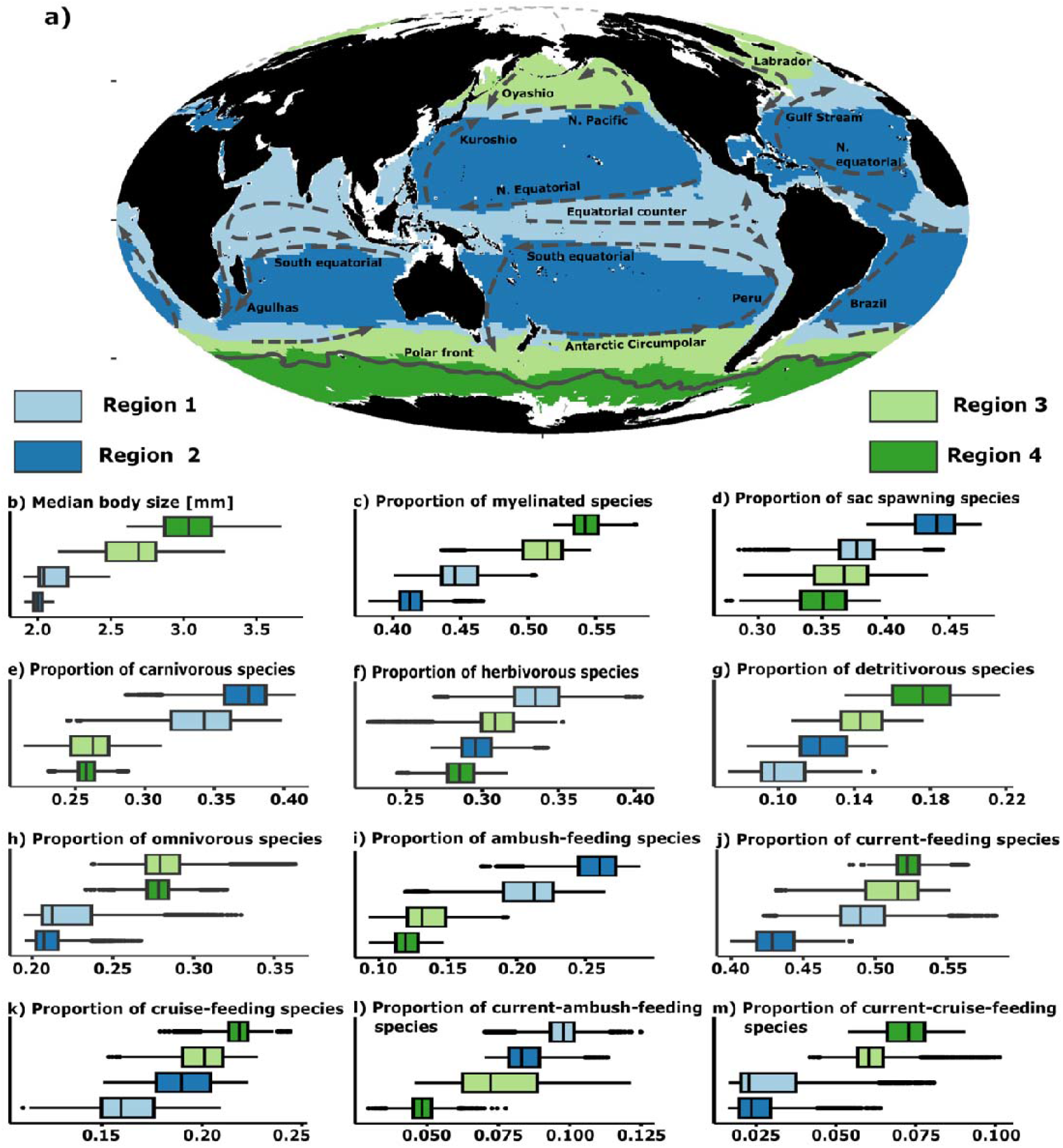
Partitioning of the global surface ocean into a) four regions according to the principal components of a Principal Component Analysis (PCA) based on the CWM trait values (Mollweide projection). Major oceanographic circulation features were overlayed to highlight their overlap between the regions’ boundaries. The distribution of CWM trait values between the four regions are shown through the boxplots: b) median body size, CWM values of c) myelinated species, d) sac spawning species, e) carnivores, f) herbivores, g) detritivores, h) omnivores, i) ambush-feeders, j) current-feeders, k) cruise-feeders, l) current-ambush-feeders, and m) current-cruise-feeders. The lower, middle, and upper boundaries of the boxplots correspond to the 25th, 50th, and 75th percentiles respectively. The lower and upper whiskers extend no further than 1.5*IQR (interquartile range) from the lower and upper hinges.

Region 1 fell within the equatorial band and comprised coastal upwelling regions and the main oxygen minimum zones. The copepod communities of Region 1 displayed lower CWM body size (median ± IQR = 2.043 mm ± 0.194), higher proportions of carnivores (0.342 ± 0.040) and herbivores (0.335 ± 0.029) but lower proportions of omnivores (0.213 ± 0.030) and detritivores (0.098 ± 0.023). The CWM values of myelinated (0.446 ± 0.026), sac-spawning (0.377 ± 0.026), and ambush-feeding (0.212 ± 0.036) were lower than in Region 2 and higher than in Regions 3 and 4 (Appendix S13).

Region 2 comprised the tropical gyres. We found that communities in Region 2 showed the lowest CW median body size (2.002 ± 0.057 mm) but the highest CWM values of ambush-feeding (0.259 ± 0.027) carnivorous (0.372 ± 0.028), and sac-spawning (0.440 ± 0.030) copepods. Conversely, the CWM values of myelinated (0.413 ± 0.016) species in Region 2 were the lowest across all four regions.

Region 3 was located poleward to the previous two regions and covered the North Atlantic, the North Pacific and the waters located between the Polar Front and the Antarctic Circumpolar Current. The CWM body size (2.691 ± 0.342 mm), the CWM values of myelinated (0.514 ± 0.028) and omnivorous (0.279 ± 0.021) species were higher there than in Regions 1 and 2. In contrast, the CWM values of sac-spawners (0.368 ± 0.041), carnivores (0.266 ± 0.026), and ambush feeders (0.132 ± 0.027) were lower.

Region 4 mainly corresponded to the Southern Ocean (i.e., grid cells south of the Antarctic Polar Front, Fig. 5a). Region 4 displayed the highest CWM body size (3.035 ± 0.321 mm), higher CWM values of myelinated (0.541 ± 0.018), omnivorous (0.278 ± 0.013), and detritivorous (0.176 ± 0.030) species, and the lowest CWM values of sac-spawners (0.351 ± 0.035). The CWM values of current-feeding (0.522 ± 0.015) and cruise-feeding (0.218 ± 0.009) species were higher than those of ambush-feeding (0.120 ± 0.016).

## 4. Discussion

### 4.1. Towards meaningful global copepod FGs in marine ecology and ecosystem modelling

We identified eleven copepod FGs based on combinations of species-level functional traits. Nine of these eleven FGs had also been found, or were nested within larger groups, in previous studies based on regional species pools (Table 1), which suggests that most of the functions performed by copepods on regional scales can also be found in the upper hundreds of meters of the global ocean.

**Table 1:** Comparison of the copepod functional groups (FGs) defined in the present study and those found in previous studies based on species and functional trait composition. The range (median ± IQR) of the species body size is given in brackets for our FGs. List of acronyms describing the functional traits: *Sma* = Small; *Lar* = Large; *Mye* = Myelinated; *Amye* = Amyelinated; *Sac* = Sac-spawning; *Cru* = Cruise-feeding; *Cur* = Current-feeding; *Amb* = Ambush-feeding; *Mix* = Mixed-feeding; *Omni* = Omnivores; *Herb* = Herbivores; *Carn* = Carnivores; *Detr* = Detritivores.

| This study | Main clade | Group composition | Pomerleau et al. (2015) - North Pacific Ocean | Benedetti et al. (2016) - Mediterranean Sea | Benedetti et al. (2018) - Mediterranean Sea | Becker et al. South Atlantic |
| --- | --- | --- | --- | --- | --- | --- |
| FG1 | *Clausocalanus* | Mye, Cru, Omni-Herb ($1.10 \pm 0.52$) | Sma, Cur+Cru, Omni-Herb (Group 6) | Sma, Cru, Omni-Herb (subset of Group 6) | Sma, Cru, Omni-Herb (Group 7) | Sma+Lar, Cru+Cur, Omni (subset of Group 6) |
| FG2 | *Oncaeidae* | Amye, Cru, Detr ($0.80 \pm 0.30$) | No equivalent | Sma, Cru, Omni-Detr (subset of Group 6) | Sma, Sac, Detr (subset of Group 5) | Sma, Amye, Cru, Carn (subset of Group 6) |
| FG3 | *Spinocalanus*, *Scaphocalanus*, *Metridia* | Mye, Mix or Cru, Detr ($2.61 \pm 1.70$) | No equivalent | Sma, Cru, Omni-Detr (subset of Group 6) | Sma, Sac, Detr (subset of Group 5) | No equivalent |
| FG4 | *Corycaeidae* | Amye, Amb, Carn ($1.22 \pm 0.70$) | No equivalent | Sma, Amb, Carn (Group 2) | Sma, Amb, Carn (Group 2) | Sma, Amye, Cru, Carn (subset of Group 6) |
| FG5 | *Sapphirinidae*, *Euchaetidae* | Cur or Cru, Carn ($4.58 \pm 2.75$) | No equivalent | Lar, Cru, Carn (Group 1) | Lar, Cru or Cur, Carn (Group 1) | Sma+Lar, Mye, Cur, Omni (subset of Group 6) |
| FG6 | *Calanus*, *Eucalanus* | Mye, Cur, Omni-Herb ($6.80 \pm 2.15$) | No equivalent | No equivalent | No equivalent | No equivalent |
| FG7 | *Calanus*, *Paracalanus*, *Calocalanus* | Mye, Cur, Omni-Herb ($1.80 \pm 2.26$) | Sma+Lar, Cur, Omni-Herb (Group 5c) | Sma+Lar, Cur, Omni-Herb (Group 4) | Sma+Lar, Cur, Omni-Herb (Groups 3 and 4) | Lar, Mye, Omni-Herb (subset of Group 6) |
| FG8 | *Candacia*, *Haloptilus*, *Heterorhabdus* | Amy, Amb or Cur, Carn ($3.40 \pm 1.97$) | No equivalent | Lar, Cru, Carn (Group 1) | Lar, Cru or Cur, Carn (Group 1) | No equivalent |
| FG9 | *Pleuromamma*, *Gaetanus*, *Labidocera* | Amy, Cur, Omni ($2.76 \pm 1.98$) | No equivalent | Sma+Lar, Cur, Omni-Herb (subset of Group 4) | Sma+Lar, Cur, Omni-Herb (subset of Group 4) | No equivalent |
| FG10 | *Oithonidae* | Amy, Amb, Omni ($1.10 \pm 0.52$) | No equivalent | Sma, Amb, Omni (Group 5) | Sma, Amb, Omni (Group 6) | Sma, Amye, Amb, Omni (A) |
| FG11 | *Acartia*, *Centropages* | Amy, Mix, Omni ($1.50 \pm 0.67$) | Sma, Amb, Omni (subset of Group 4) | Sma, Mix, Omni (Group 3) | Sma, Mix, Omni (subset of Group 4) | Sma, Amye, Amb, Omni (A) |

Only the new FG3 and FG6 have no counterparts in previous studies. Their most suitable habitats lie towards the poles (Fig. 4), whose copepod communities were not covered by the regional studies mentioned. The copepods of FG3 (*Spinocalanus* spp., *Metridia* spp., or *Scaphocalanus* spp.) are known to mainly inhabit deeper ocean layers where they feed on sinking particulate organic matter and zooplankton carcasses (Yamaguchi et al., 2002; Sano et al., 2013). Therefore, this group contributes to the remineralization of organic matter and marine snow at higher latitudes and/or in colder conditions. FG6 comprises the largest (6.80 ± 2.15 mm) current-feeding omnivorous-herbivorous copepods from the Calanidae (e.g., *Calanus hyperboreus, C. glacialis, Eucalanus bungii* or *Rhincalanus gigas*). Many of these large copepods are not present in the warm and oligotrophic conditions of the Mediterranean Sea, hence their absence in Benedetti, Gasparini, & Ayata (2016) and Benedetti, Vogt et al. (2018). FG6 is a key group for the biological carbon pump (Jónasdóttir et al., 2015; Visser et al., 2017; Steinberg & Landry, 2017), as it represents large-bodied grazers that can actively feed on microphytoplankton, perform relatively strong vertical migrations and generate large, fast sinking pellets (Stamieszkin et al., 2015; Ohman & Romagnan 2016). This is confirmed by the position of FG6 in niche space (Fig. 3) as it is affiliated with turbulent, seasonally varying conditions with higher nutrient and chlorophyll-a concentrations. The species of FG7 fill similar functions but display smaller body sizes (1.80 ± 2.26mm; e.g., *C. helgolandicus, C. pacificus* and the Paracalanidae), hence we expect their contribution to size-mediated functions to be lower. The ecological roles and functions of the other nine FGs have already been discussed in previous studies, and our results support the description of their ecological roles. Here, we found a larger range of FGs compared to previous studies due to three main reasons: (i) we investigated a global pool of species (hundreds of species instead of tens), (ii) we accounted for myelination as an additional trait contrary to Benedetti, Gasparini, & Ayata (2016) and Benedetti, Vogt et al. (2018), and (iii) we retained body size as a continuous trait instead of using size classes.

Overall, the largest differences in niche space (Appendix S14) occurred between two main sets of FGs: (i) myelinated free-spawning current-feeding herbivores of various sizes (FG3, 6, 7 and 9), and (ii) amyelinated sac-spawning small-bodied, or large-bodied, cruise- and ambush-feeding detritivores and carnivores (FG2, 4 and 5). The first set is associated with conditions of stronger mixing and higher nutrient and chlorophyll-a concentrations (Appendix S14). The species in these FGs also have larger niche widths, i.e. they display broader tolerances to variations in environmental conditions. Therefore, these FGs are found more frequently in high latitude environments, or boundary current systems, where either seasonality or physical mixing lead to higher mean annual productivity (Sarmiento & Gruber, 2006; Roy 2018). In contrast, the FGs of the second set display narrower niches and are associated with conditions typical of the warmer tropical oligotrophic gyres (i.e., weaker water mixing and lower nutrient and chlorophyll-a concentrations; Appendix S14). Meanwhile, the FGs that are in the center of the niche space (FG1, 8, 10 and 11) could not be associated to any particular environment, likely because they include genera with a wide latitudinal distributions.

FGs can also combine species that are known to inhabit different vertical layers of the water column (e.g., FG9 which gathers epipelagic species with meso-bathypelagic ones). This is because depth preferences cannot be considered as a functional trait per se (Litchman et al., 2013). Rather, the depth range of a species reflects the outcome of the interactions between its traits and the abiotic and biotic conditions. Therefore, our approach likely overlooks vertical habitat partitioning that occur between or within FGs.

The continuum of FGs in niche space was also found in functional trait space. Indeed, the FGs of sets (i) and (ii) were often found on opposite sides of the FAMD dimensions (Appendix S12). The global distribution of the FGs and CWM trait values further support the statements above (Figs. 3 and 4): larger myelinated free-spawning and current- and cruise-feeding copepods occur more frequently near the poles, whereas smaller amyelinated sac-spawning and ambush-feeding copepods tend to occur in tropical oligotrophic gyres. Similar continuums of copepod functional traits and niches were found at regional (Table 1) to global scales (Brun et al., 2016).

In summary, our results support the view that planktonic copepods display a continuum of functional traits with a strong latitudinal gradient (Figs. 4 and 5; Appendix S9), driven by global gradients in abiotic conditions. Consequently, such a continuum should be explicitly represented in marine ecosystem models (e.g. similar to Serra-Pompei et al., 2020), which too often rely on a few size classes only (Le Quéré et al., 2005). These need to include FGs that are both distinct, and important in terms of their biomass. To rank the present FGs based on their contribution to total copepod community abundance in the upper ocean, we implemented a preliminary synthesis of copepod abundance observations from various data sources (Appendix S16). Examining the relative contribution of the eleven FGs to mean annual community abundance in the upper 200m (Fig. S16b) showed that three FGs emerge as the most abundant, regardless of latitude and sampling gears: FG10 (Oithonids; small ambush-feeding omnivores), FG2 (Oncaeids; small cruise-feeding detritivores) and FG1 (*Clausocalanus* spp.; small cruise-feeding omnivores-herbivores). These FGs are thus likely key the development of future ecosystem models. Future work is needed to better assess the contribution of the copepod FGs to biomass and clarify their relative priorities for inclusion in ecosystem models.

### 4.2. Why do copepod functional traits show diverging biogeographic patterns in the ocean?

The largest dissimilarity in community trait expression was found between regions 2 and 4 (Fig. 5b-m) and was driven by differences in CWM body size, myelination, carnivory and ambush-feeding vs. current-feeding (Appendices S9, S10 and S15). We found the boundaries of the four main regions to overlap with the trajectories of well-known boundary currents as well as equatorial counter currents and fronts (Fig. 5). Trait patterns emerge from environment-driven distribution models, so they reflect the fact that certain trait combinations are more competitive than others under varying temperature and food conditions (Barton et al., 2013; van Someren Gréve et al., 2017; McGinty et al., 2021).

Median copepod body size decreased from the poles to the equator with a slight increase in upwelling systems. Such a pattern is primarily driven by the strong negative relationship between the body size of marine ectotherms and temperature, according to Bergmann’s rule (Brun et al., 2016; McGinty et al., 2018; Evans et al., 2020; Brandão et al., 2021; Campbell et al., 2021). The processes underlying Bergmann’s rule remain debated, but it is likely that warmer temperatures (or a factor confounded with temperature) decrease growth efficiency and/or promote the maturation of adults at smaller body sizes (Atkinson 1994; Isla et al., 2008).

Bergmann’s rule occurs at the intraspecific and interspecific levels, meaning that warming-induced decreases in median body size can emerge if clades of small species replace large-bodied ones along a latitudinal gradient. Such a turnover in size classes has been observed as well (Evans et al., 2020; Brandão et al., 2021) and is supported by our SDM projections. Indeed, functional traits that represent shifts in clade composition such as myelination, spawning strategy, carnivory or feeding modes display latitudinal gradients that are positively or negatively correlated with the body size gradient (Appendices S9 and S10). The latitudinal patterns of these functional traits likely result from changes in food availability, quality, and predation pressure (Horne et al., 2016; van Someren Gréve et al., 2017; Roy, 2018).

Lipid-rich myelin sheaths enable a faster conduction of nerve responses. This promotes faster reaction times and thus more efficient feeding or escape behaviors (Lenz, 2012). Myelin sheaths are cholesterol-rich so they require larger metabolic investments of dietary lipids (Lenz, 2012), which helps explain why copepod communities show larger proportion of small amyelinated taxa in tropical gyres where smaller, lipid-poor phytoplankton dominate (Roy, 2018). Therefore, the proportion of myelinated species follows the same spatial pattern as body size and current-feeding (Fig. 5b,c,j,m) and peaks in productive environments characterized by larger and lipid-rich plankton (Roy 2018).

Similarly, sac-spawning is an energy-conservative spawning strategy, as it reduces egg-mortality at the cost of fecundity and hatching speed. The increased proportions of sac-spawners in tropical gyres (Fig. 5d) could reflect an adaptation to limited food availability (Kiørboe & Sabatini, 1994; Barton et al., 2013) and higher rates of carnivory (Fig. 5e; Woodd-Walker et al., 2002) and egg cannibalism among copepods (Ohman & Hirche, 2001; Segers & Taborsky, 2011). Ambush-feeding is a passive feeding mode that lowers predation risk and energy costs compared to current- and cruise-feeding, but at the expense of feeding efficiency (Kiørboe, 2011a; van Someren Gréve et al., 2017). Consequently, the increased proportion of ambush-feeders in region 2 (Fig. 5i) may result from trade-offs in functional traits expression driven by abiotic and biotic filtering (Litchman et al., 2013), as oligotrophic gyres seem to promote food-webs with increased carnivorous predation (Fig. 5e) and resource competition (Woodd-Walker et al., 2002; Prowe, Visser, Andersen, Chiba, & Kiørboe, 2019). Since our approach is based on presence data and habitat suitability indices rather than abundances, it likely underestimates the contribution of very abundant ambush-feeding species like *Oithona similis* at high latitudes (Pinkerton et al., 2010; Prowe et al., 2019; Becker et al., 2021). As a result, we probably underestimate the proportion of ambush-feeders in regions such as the Southern Ocean compared to Prowe et al. (2019), although these authors discarded other ambush-feeding copepods such as the Corycaeidae (Benedetti et al., 2016; Brun et al., 2017). These elements support the fact that our modelled copepods functional traits patterns emerge from interactions between environmental conditions and the relative fitness resulting from different trait combinations.

### 4.3. Caveats and conclusion

Our results are sensitive to: (i) the way the species were positioned in a functional space and how the FGs were defined from the latter, (ii) the quantity and quality of the functional trait data considered, and (iii) the SDMs chosen to generate the CWM traits values. We investigated the sensitivity of our FGs to the clustering approach (Appendix S4). We assessed how using this alternative dimension reduction analysis affected the quality of the functional trait space by computing the Area Under the Curve (AUC) criterion (Mouillot et al., 2021). We found an AUC value of 0.81, which is substantially higher than the recommended 0.7 threshold and indicates a “high quality trait space”. Plus, all the functional dendrograms emerging from alternative clustering approaches showed fairly similar structure (mean Baker’s Gamma correlation coefficient was 0.75 ± 0.12), indicating that they all lead to similar FGs. The traits chosen for this study only cover a fraction of the traits mentioned in the literature (Litchman et al., 2013; Appendix S3). Therefore, our study likely underestimates the true diversity of copepod functions in the ocean. More field observations and lab experiments are required to document the existing traits of many copepod species (Barton et al., 2013).

We found some regional differences between the mean annual HSI projections of the SDMs. The GLM estimated higher mean HSI than the two other model types in the Southern Ocean (Appendix S8). Such variability stems from how the various SDMs cope with limited predictors and species data availability in winter conditions at the very high latitudes. Here, the less complex response curves of GLMs (Merow et al., 2014) lead to higher average HSI in very cold temperatures compared to GAMs. The HSI patterns obtained from the GAMs and ANNs were closer to the latitudinal diversity gradients that were previously observed for marine ectotherms (Tittensor et al., 2010; Benedetti et al., 2021). Interestingly, inter-SDMs variability was much lower when looking at CWM traits projections rather than HSI patterns (Appendix S8). The fact that CWM body size and CWM myelination show very similar spatial patterns and ranges to the estimates of Brun et al. (2016) gives us further confidence in our spatial projections.

To conclude, we recommend the inclusion of multiple FGs of copepods in marine ecosystem models, based on our functional dendrogram (Fig. 2), with the level of complexity of copepod representation reflecting not only the FGs dominating community biomass at the scale of the study region(s), but also the traits of interest. Ideally, future global marine ecosystems, that cannot efficiently include 11 copepod groups, will include a few (3-4) FGs that cover the main gradients observed in trait space (Fig. 2; Appendix S12), niche space (Fig. 3), geographical space (Figs. 4, 5; Appendices S9, S10) and that represent the majority of global or regional biomass. For other ecological applications or to represent food-web interactions across trophic levels (Serra-Pompei et al., 2020), these groups could be split into further FGs with specific characteristics (e.g., size classes, high lipid content, grazing and mortality rates etc.). Future studies are required to improve the coverage of functional trait data to include other taxonomic groups and more quantitative traits (Appendix S3). Ongoing compilations of zooplankton biomass data will also help rank the regional to global importance of the present FGs, and to assess links between ecosystem function and service provision.

## Acknowledgements

This project has received funding from the European Union’s Horizon 2020 research and innovation program under grant agreement No. 862923. This output reflects only the author’s view and the European Union cannot be held responsible for any use that may be made of the information contained therein. We thank all contributors involved in the plankton species field sampling and identification throughout the world and we acknowledge the efforts made to deposit the data on publicly available online archives. We thank Luke Gregor for editing the language of an early version of the manuscript. We thank Dr. Maria Grazia Mazzocchi and an anonymous reviewer for their constructive comments. No permits were needed to carry out the present study.

## Data Availability Statement

The copepod species occurrences data used to train the species distribution models are publicly available on Zenodo (https://doi.org/10.5281/zenodo.5101349). The species functional traits table is available as Supplementary Appendix S2. All R codes are accessible on the GitHub account of J.W. (https://github.com/jonas-wydler) and are also available on Zenodo (https://doi.org/10.5281/zenodo.7050567).

## Biosketch

**Fabio Benedetti** is a postdoctoral researcher and **Meike Vogt** a senior research scientist in the Environmental Physics (UP) group of ETH Zürich. Both share broad interests in plankton biogeography and functional diversity and theirs links with biodiversity, ecosystem function and biogeochemical cycles in the global ocean. F.B. is a macroecologist specialized in trait-based approaches and plankton diversity modelling. M.V. is a marine ecosystem modeler specialized plankton functional types. **Jonas Wydler** has successfully completed his MSc degree in Environmental System Sciences at ETH Zürich, under the supervision of F.B. and M.V., which is the subject of this work.

F.B. and M.V. co-designed the study and F.B. collated the data used in the analyses and provided expertise with regard to every methodology used. J.W. conducted the numerical analyses and wrote the first version of the manuscript under the supervision of F.B. and M.V. F.B. wrote the final version of the manuscript with input from both M.V. and J.W.

